# Differences in the dynamics of the tandem-SH2 modules of the Syk and ZAP-70 tyrosine kinases

**DOI:** 10.1101/2021.07.20.453126

**Authors:** Helen T Hobbs, Neel H Shah, Jean M Badroos, Christine L Gee, Susan Marqusee, John Kuriyan

**Affiliations:** Department of Chemistry, University of California, Berkeley, United States; California Institute for Quantitative Biosciences, University of California, Berkeley, United States; Howard Hughes Medical Institute, University of California, Berkeley, United States; Department of Molecular and Cell Biology, University of California, Berkeley, United States; Molecular Biophysics and Integrated Bioimaging Division, Lawrence Berkeley National Laboratory, Berkeley, United States; Department of Chemistry, Columbia University, 3000 Broadway, New York, NY 10027

**Keywords:** protein dynamics, hydrogen-deuterium exchange mass spectrometry, evolution, tyrosine kinase signaling, conformational landscape

## Abstract

The catalytic activity of Syk-family tyrosine kinases is regulated by a tandem-SH2 module (tSH2 module). In the autoinhibited state, this module adopts a conformation which stabilizes an inactive conformation of the kinase domain. The binding of the tSH2 module to doubly-phosphorylated tyrosine-containing motifs necessitates a conformational change, thereby relieving kinase inhibition and promoting activation. We determined the crystal structure of the isolated tSH2 module of Syk and find, in contrast to ZAP-70, that its conformation more closely resembles that of the peptide-bound state, rather than the autoinhibited state. Hydrogen-deuterium exchange by mass spectrometry, as well as molecular dynamics simulations, reveal that the dynamics of the tSH2 modules of Syk and ZAP-70 differ, with most of these differences occurring in the C-terminal SH2 domain. Our data suggest that the conformational landscapes of the tSH2 modules in Syk and ZAP-70 have been tuned differently, such that the auto-inhibited conformation of the Syk tSH2 module is less stable. This feature of Syk likely contributes to its ability to more readily escape autoinhibition when compared to ZAP-70, consistent with tighter control of downstream signaling pathways in T cells.

## INTRODUCTION

The adaptive immune system facilitates highly specific and robust responses to a wide variety of antigens. The primary drivers of this complex system are lymphocytes, known as B and T cells.^1^ Together, they work to mount responses to antigens, as well as to store memories of those antigens. B cells are primarily involved in humoral immunity through the production and secretion of antibodies, while T cells play various roles in cell-mediated immunity.^1,2^ This intricate system arose approximately 450 million years ago in a common ancestor of jawed vertebrates.^3,4^ Two genome-wide duplication events resulted in the acquisition of many paralogous genes with essential roles in B and T cell-mediated immunity.^3^ These genes encode important proteins found in signal-transduction pathways that are activated following the engagement of cell-surface antigen receptors, such as the B cell receptor (BCR) or T cell receptor (TCR). The regulation of these signaling proteins is critical for B and T cell-mediated immunity.

One such set of paralogous genes encodes the two Syk-family kinases, spleen tyrosine kinase (Syk) and zeta-chain-associated protein kinase 70 (ZAP-70).^5^ Syk and ZAP-70 are critical triggers of early signaling events in B cells and T cells respectively. In B cells, following BCR engagement, the Src-family kinase Lyn phosphorylates the cytoplasmic tails of the BCR on tyrosine residues located in immunoreceptor tyrosine-based activation motifs (ITAMs) (Figure 1A, left).^6,7^ Syk is recruited to these phosphorylated ITAMs through its tandem-SH2 module (tSH2 module) and is activated by ITAM binding. In T-cells, the Src family kinase Lck phosphorylates the ITAM-containing cytoplasmic tails associated with the TCR leading to recruitment and activation of ZAP-70 (Figure 1A, right).^8^ Syk differs from ZAP-70 in that it can directly phosphorylate ITAMs and also phosphorylates itself on the kinase activation loop, further increasing activity.^9,10^ While not required, the activity of Lyn has been shown to increase BCR sensitivity to monomeric antigens and to mediate more rapid signaling than what would occur from Syk activity alone.^11^ In contrast, ZAP-70 requires a Src-family kinase for activation, reflecting stricter regulation of the activation of T cells.

**Figure 1.**
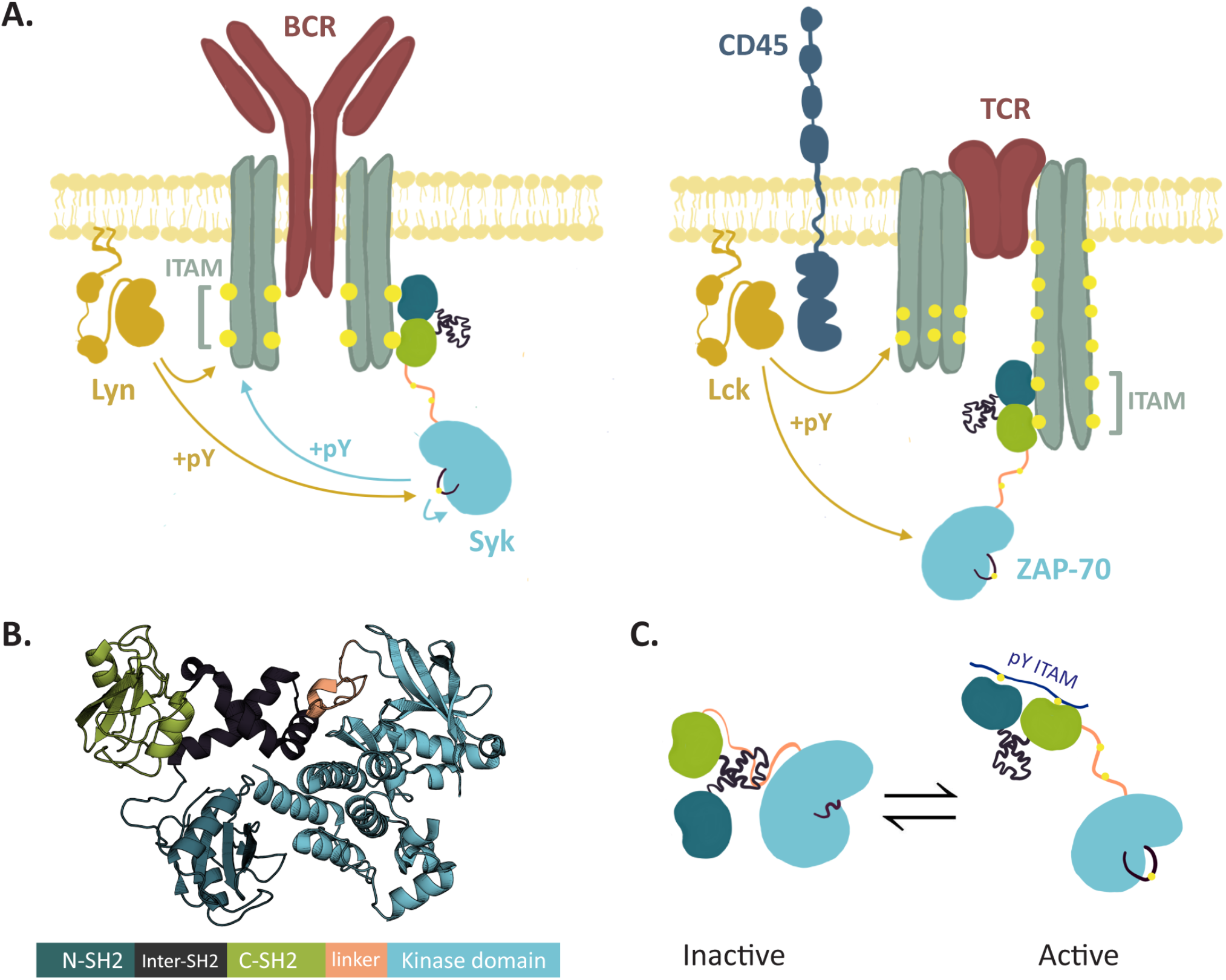
Structure and function of the Syk family kinases. **A**. Early T cell signaling events, left. Early B cell signaling events, right. In B cells and T cells a Src family kinase (yellow) phosphorylates ITAMs (phsophotyrosine denoted by yellow spheres) near the receptor. Binding of the tSH2 module to phosphorylated ITAMs recruits the kinase to the receptor. ZAP-70 signaling requires Lck, the Src family kinase, but Syk can initiate some signaling, even in the absence of Lyn, the B cell Src family kinase. **B**. Domain architecture and autoinhibited structure of Syk family kinases (PDB: 4FL2, autoinhibited human Syk) **C**. Activation of Syk family kinases by conformational change in the tSH2 module and phosphorylation. The tSH2 module binds to a doubly-phosphorylated ITAM (blue) and phosphorylation of tSH2-kinase linker stabilizes the open, active conformation.

Syk and ZAP-70 share a common domain architecture (Figure 1B), which includes two Src-homology 2 (SH2) domains connected by a helical linker, comprising the tandem-SH2 module (tSH2 module). A flexible linker (SH2-kinase linker) connects the tSH2 module to the kinase domain. Like most protein kinases, Syk and ZAP-70 can switch from an autoinhibited conformation (“OFF state”) to a catalytically-competent conformation (“ON state”).^12–14^ As an example, the OFF state for Syk is depicted in Figure 1B (PDB: 4FL2).^12^ In this conformation, the docking of the SH2-kinase linker on the face of the kinase domain distal to the active site impedes the inward rotation of the αC-helix.^14^ The rotation of the αC-helix is key to the conformational transition between the ON and OFF states and is coupled to the conformation of the activation loop at the active site cleft.^15^ In particular, the sidechain of Tyr 352 in the SH2-kinase linker of human Syk (Tyr 319 in human ZAP-70) is wedged in the interface between the αC-helix and the N-lobe of the kinase domain, stabilizing the OFF state.^13,14^ This state is further stabilized by the tSH2 module, which adopts a conformation in which the relative orientation of the two SH2 domains is inconsistent with ITAM binding.^16,17^ ITAM binding triggers a conformational change towards the ON state, which is further enforced by phosphorylation of the SH2-kinase linker at Tyr 348 and Tyr 352 (Tyr 315 and Tyr 319 in ZAP-70) (Figure 1C).^12,13^ The crystal structure of the ITAM-bound tSH2 module has been reported for both Syk (PDB:1A81) and ZAP-70 (PDB:1M61).^16,17^ Overall, these structures are very similar. For ZAP-70, the crystal structure of the ITAM-free tSH2 module has also been reported. In this structure the C-terminal SH2 (C-SH2) domain is rotated substantially from its position in the ITAM-bound structure.^18^ This conformation is similar to that observed in autoinhibited ZAP-70.^12^

We now present the crystal structure of the isolated, ITAM-free tSH2 module of Syk, which has not been previously reported. Unlike ZAP-70, the Syk tSH2 module crystallizes in a conformation similar to that in the ITAM-bound structure and distinct from its conformation in the autoinhibited structure. To further understand how the tSH2 module differs between Syk and ZAP-70, we studied the dynamics of these proteins using hydrogen-deuterium exchange mass spectrometry (HDX-MS) and also ran all-atom molecular dynamics simulations. The resulting data are in agreement with previous studies that showed that the C-SH2 of ZAP-70 undergoes a large conformational change upon binding of an ITAM and that it is dynamic in the absence of a bound peptide.^19,20^ Conversely, we find that the C-SH2 of Syk is relatively rigid, even in the unbound state, and does not undergo a similar conformational rearrangement upon binding a peptide. The bias of the Syk tSH2 module towards a conformation resembling that found in the ON state of the kinase, even in the absence of ITAM peptides, suggests that the auto-inhibited state of Syk is less stable than that of ZAP-70.

## RESULTS

### Crystal structure of the ITAM-free Syk tSH2 module

We determined the structure of the ITAM-free Syk tSH2 module at 3.2 Å resolution by x-ray crystallography. The structure was solved by molecular replacement using Chain F from the ITAM-bound structure of Syk (PDB: 1A81).^17^ The crystal conditions, space group, and unit cell dimensions, described in Table S1, differ from those reported for the ITAM-bound structure.^17^ Additionally, while some of the crystal packing interfaces are similar to those of the ITAM-bound complex, not all of the interfaces are the same, suggesting that similarity in conformation is not due to crystal packing. As in the structure of the ITAM-bound Syk tSH2 module (PDB:1A81), we observe a small degree of conformational heterogeneity in the six molecules in the asymmetric unit. When all six tSH2 modules are aligned on their N-SH2 domains, as in Figure 2A, the positions of the C-SH2 domains and the inter-SH2 linker are slightly variable. The most extreme differences between molecules in the asymmetric unit, represented by the darkest and lightest red structures in Figure 2A, results in a displacement of the C-SH2 of 2.7 Å.

**Figure 2.**
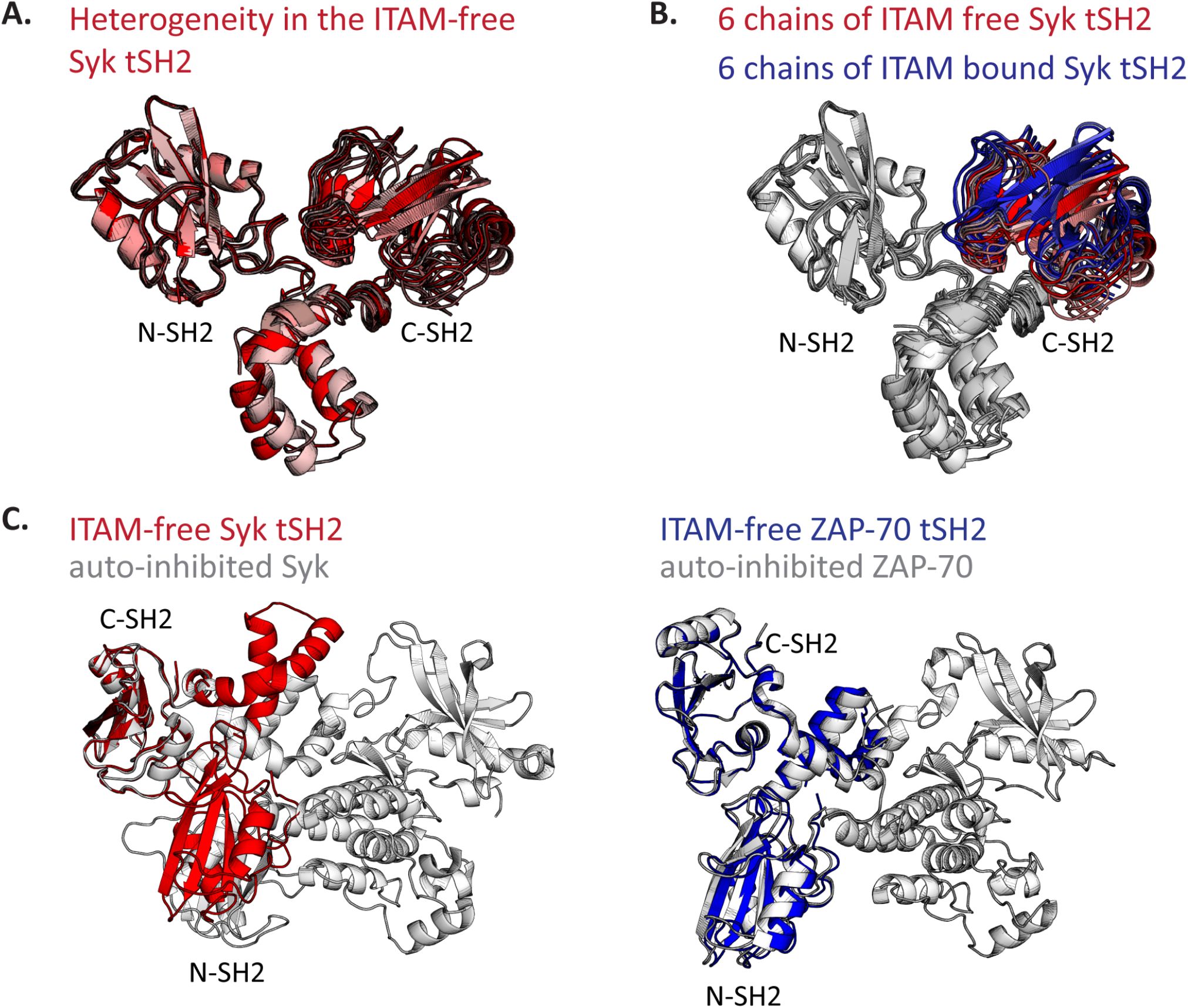
The crystal structure of the ITAM-free tSH2 module is similar to that of the ITAM- bound Syk tSH2 module. **A**. The six molecules of ITAM-free Syk tSH2 aligned on the N-SH2 domain, showing the observed conformational heterogeneity. **B**. Alignment on N-SH2 of all molecules in the ITAM-bound structure (PDB: 1A81, blue) and all in the ITAM-free structure (red). Only the C-SH2 is colored. **C**. Left, alignment on the C-SH2 domain of the isolated, ITAM-free Syk tSH2 module (red) to that in the full-length auto-inhibited Syk structure (PDB:4FL2, white), resulting in a ∼36 Å displacement of the N-SH2 domain of the ITAM-free tSH2 module with respect to its position in the auto-inhibited state. Right, the ITAM-free ZAP-70 tSH2 module (PDB: 1M61, blue) aligns well to the full-length auto-inhibited structure of ZAP-70 (PDB:4K2R, white), right.

We find that, unlike the ZAP-70 tSH2 module, the ITAM-free Syk tSH2 module adopts a conformation remarkably similar to that observed in the ITAM complex and distinct from the conformation observed in the autoinhibited kinase structure (Figure 2B and C). When the molecules from the ITAM-bound structure (PDB:1A81) and the molecules from the ITAM-free structure are aligned on their N-SH2 domain, the displacement of the C-SH2 domain ranges between ∼0.8 and 6.6 Å (value varies based on the molecules used in the alignment). Overall, the tSH2 modules in our ITAM-free structure represent a set of conformations in which the SH2 domains have moved apart, with respect to the ITAM-bound structure (Figure 2B). When an alignment on the C-SH2 domain is performed using the tSH2 module from the autoinhibited, full-length structure of Syk (PDB:4FL2), the displacement for the C-SH2 in the ITAM-free structure is ∼30.5 Å, reflecting a large change in the conformation. The structure of the ITAM-free Syk tSH2 module described here suggests that it favors the conformation corresponding to the ON state, unlike the tSH2 module of ZAP-70, which prefers the conformation of the OFF state.

### Analysis of the dynamics of the tSH2 modules by hydrogen-deuterium exchange mass spectrometry

To further interrogate the conformational ensembles of the Syk and ZAP-70 tSH2 modules we turned to HDX-MS. This technique monitors the time course of exchange of backbone amide hydrogens with heavier deuterons in the solvent. The ability of an amide to undergo exchange is directly related to its local structure and stability.^21^ Briefly, in our experiment a protein is diluted into deuterated buffer and allowed to undergo hydrogen-deuterium exchange. After various amounts of time the exchange is quenched by lowering the pH and temperature. The quench sample is then cleaved into peptides using acid proteases, and the extent of exchange for each peptide is monitored by mass spectrometry by following the increase in molecular weight of each peptide (see Materials and Methods). We carried out HDX-MS on both the Syk and ZAP-70 tSH2 modules in the ITAM-bound and ITAM-free states. A comparison of deuterium uptake for the same peptide in different states reports on changes in the secondary structure and/or stability. For each protein state we analyzed approximately 300 individual peptides (average length of ∼11 amino acids), corresponding to 100% sequence coverage for Syk with an average redundancy (number of peptides reporting on a residue) of 11.6 and 99% sequence coverage for ZAP-70 with an average redundancy of 12.5.

The effect of ITAM binding on the deuterium uptake is depicted for a select set of peptides as a difference heatmap in Figure 3; these peptides represent the typical uptake pattern observed for all peptides in each region. These data indicate that, as expected, the amide groups in both Syk and ZAP-70 are more protected from exchange when bound to an ITAM (blue on the heatmap in Figure 3), suggesting a ligand-induced global stabilization of the entire domain, as was previously observed for the Syk tSH2 module.^22^ The extent and pattern of increased protection in Syk and ZAP-70 due to ITAM-binding differ most noticeably in regions in and around the C-SH2 domain, corresponding to residues 161-192 in Syk and 156-187 in ZAP-70. In Syk, the ITAM-free and ITAM-bound conformations show comparable exchange patterns in this region, with a possible hint of protection in the bound state. In ZAP-70, however, peptides from this region exchange rapidly in the absence of ITAM, and binding of ITAM slows the exchange, similar to what is observed in both Syk states. Together these data suggest that even in the absence of ITAM, the Syk tSH2 module adopts a conformation akin to what would be observed in the ON state of the kinase and distinct from the conformation observed in the OFF state. This tuning of the conformational landscape of Syk likely contributes to the instability of its auto-inhibited state.

**Figure 3.**
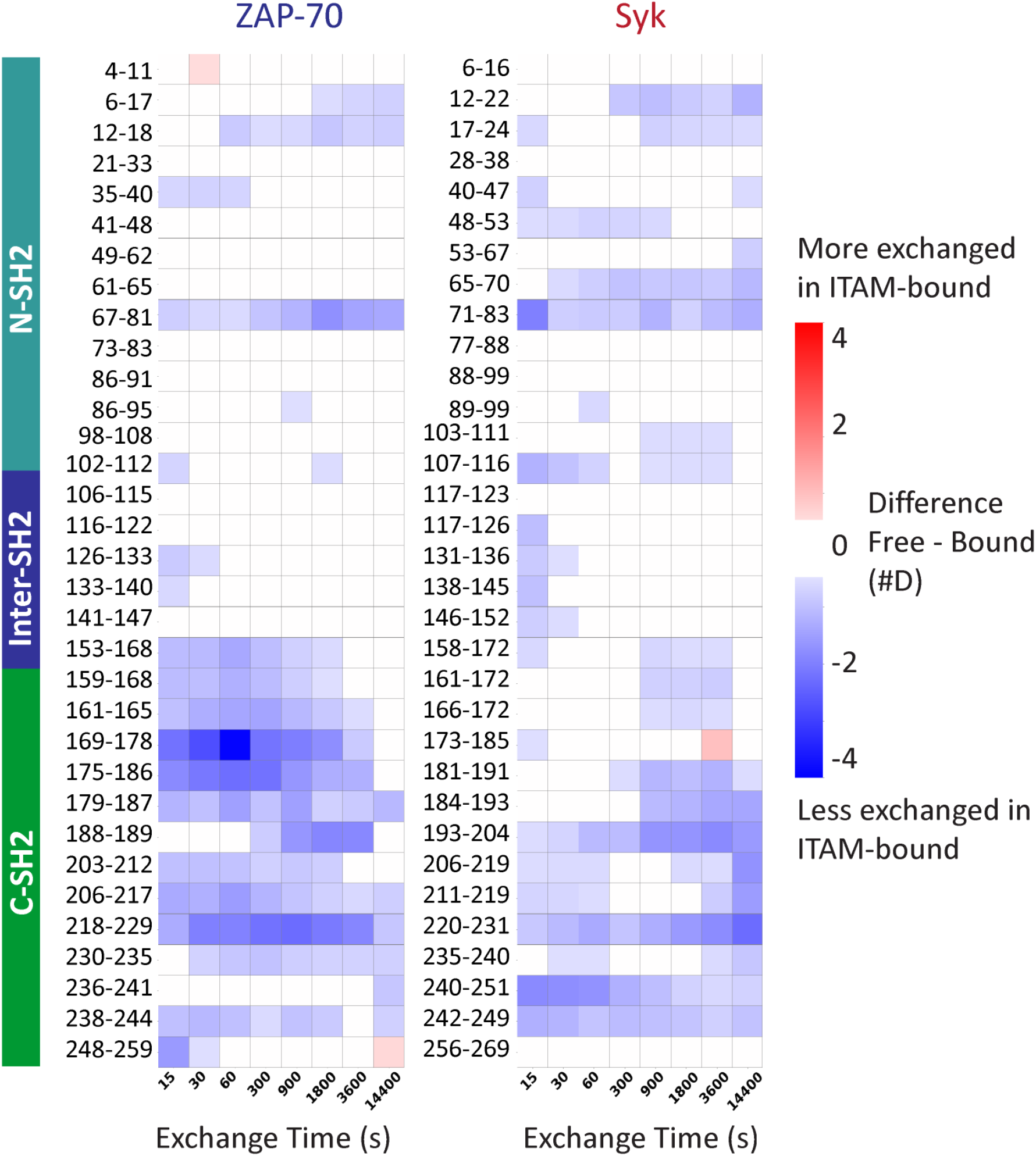
The Syk and ZAP-70 tSH2 modules show different deuterium uptake in and around the C-SH2. Heat maps from representative peptides (residue numbers noted on the left) depicting differences in deuteration between the ITAM-bound and ITAM-free states of both tSH2 modules as measured by HDX-MS (n = 3). Red squares indicate increased deuterium exchange in the ITAM-bound state and blue squares indicate decreased deuterium uptake in the ITAM-bound state. A difference of at least ±0.5 Da was considered significant, and white squares indicate no change in deuterium uptake.

**Figure 4.**
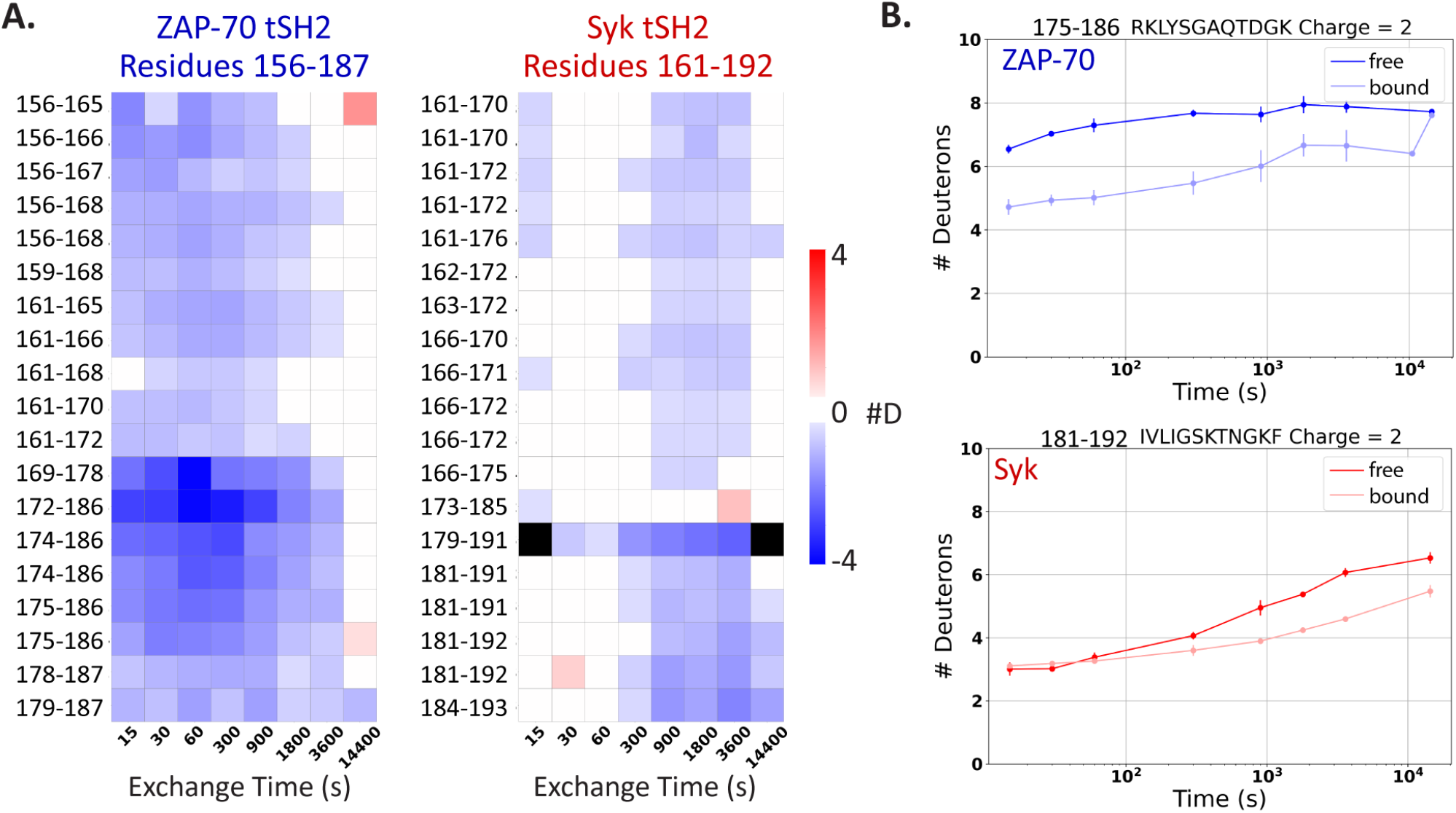
The more flexible C-SH2 domain of ZAP-70 is rigidified by ITAM binding. **A**. Deuteration difference heatmaps of all peptides, including different charge states, from the region spanning residues 161-192 in Syk (156-187 in ZAP-70). Peptides in ZAP-70 show a greater difference in deuteration between the ITAM-free and bound state. **B**. Uptake plots from similar peptide in ZAP-70 and Syk. The peptides in Syk exchange similarly in the ITAM-free (dark blue for ZAP-70 and dark red for Syk) and bound states (light blue for ZAP-70 and light red for Syk). The corresponding peptides in ZAP-70 exchange similarly to Syk in the ITAM-bound state but are more flexible in the absence of peptide.

### Molecular dynamics simulation of the tSH2 modules of Syk and ZAP-70

We ran all-atom molecular dynamics simulations starting from the ITAM-bound structures to gain insight into the differences in dynamics between the tSH2 modules of Syk and ZAP-70. After removing the ITAM, we generated five trajectories for each protein, three extending for 500 ns and two for one µs. For both Syk and ZAP-70, over the course of simulation, the N-and C-SH2 domains relax so that they are positioned farther apart than observed in the starting structure, as measured by the distance between the centers of mass of each SH2 domain (Figure 5). This movement occurs to a much greater extent in ZAP-70 than in Syk. Despite the significant separation of the SH2 domains in ZAP-70, the final structure achieved during simulation does not have the same rotation of the C-SH2 domain as observed in the ITAM-free ZAP-70 crystal structure, presumably because of the limited timescale of our simulations. In Syk, the two SH2 domains drift apart from their initial positions but still remain close together; ZAP-70 accesses a broader range of inter-SH2 distances than Syk (Figure 5). These data suggest that the tSH2 module of ZAP-70 adopts a conformation incompatible with peptide binding in the absence of ITAM.

**Figure 5.**
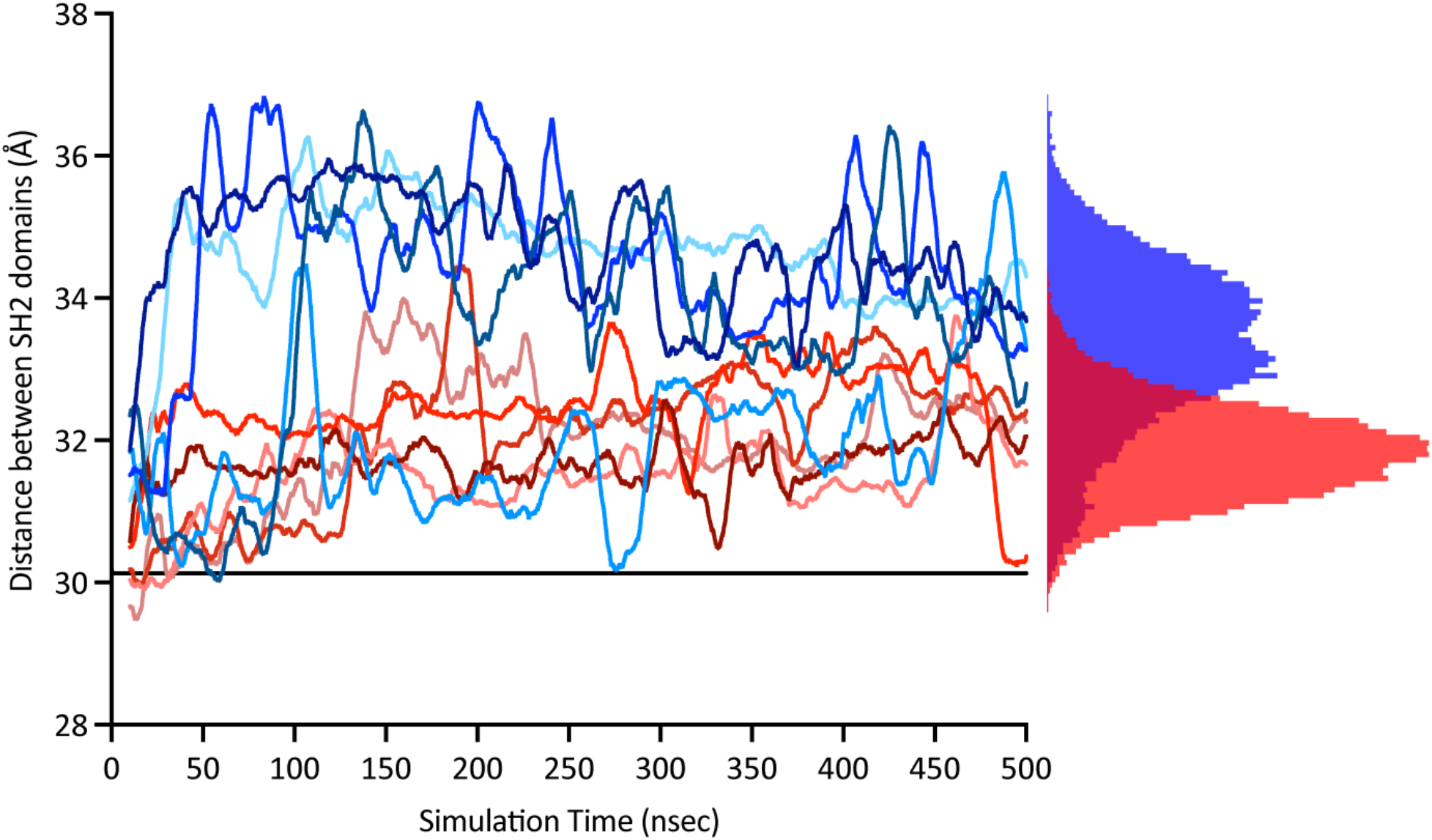
The SH2 domains of the ITAM-free ZAP-70 move apart following the removal of the bound peptide. The distance between the center of mass of the N-and C-SH2 domains in Syk tSH2 (red) and ZAP-70 tSH2 (blue). Simulations are smoothed over 10nsec. The histogram depicts the distribution of inter-SH2 distances sampled by either Syk (red) or ZAP-70 (blue).

### Sequence differences between the ZAP-70 and Syk tSH2 modules are conserved

We compiled a list of extant Syk-family kinase sequences through homology searches against databases of metazoan proteins.^23^ The organisms analyzed had zero, one, or two distinct Syk-family kinases (Figure 6A). Based on our analysis, the appearance of two Syk-family kinases in the genomes of modern organisms occurs at the transition from jawless fish to bony fish. This transition corresponds to one of the two genome-wide duplication events that correlate with the emergence of the adaptive immune system.^3^ In total, we compiled 183 sequences from 96 organisms, which were used to make a multiple sequence alignment (MSA).^24^

**Figure 6.**
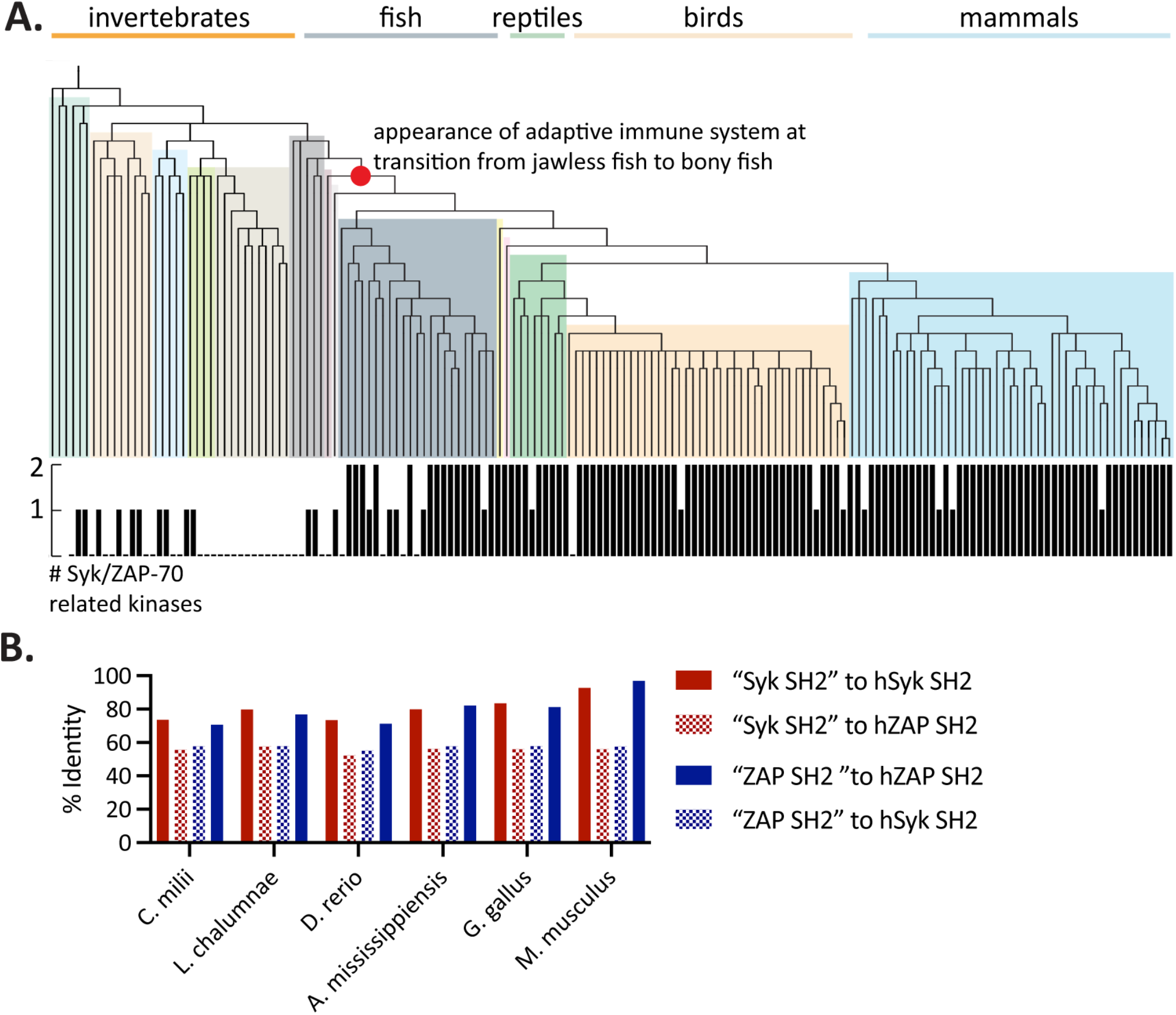
The differences between tSH2 modules are conserved in extant organisms. **A**. Species tree generated based on the taxonomic relationships between organisms searched for Syk-family kinases.30 The occurrence of Syk-family kinases in each organisms is denoted on the bar graph below the tree. The occurrence of two Syk-family kinases in the genome corresponds to the emergence of jawed vertebrates (denoted by red circle). **B**. Alignment of only the tSH2 modules to either the human Syk or human ZAP-70 SH2 modules, from a representative organism from that order (common names: shark, lungfish, zebrafish, alligator, chicken, mouse).

In those metazoans with two Syk-family kinases, each protein could reasonably be assigned as a Syk or ZAP-70 ortholog based on percent identity to the corresponding human sequence. Notably, although this sequence analysis used the full-length protein sequences, the tSH2 module sequence alone can be used to predict whether the gene corresponds to a Syk or a ZAP-70 (Figure 6B). From cartilaginous fish (*C. milii*) to mice (*M. musculus*), the tSH2 modules of Syk and ZAP-70 can be readily differentiated, with one aligning better to human Syk and the other to human ZAP-70. The tSH2 modules within a species remain approximately 55% identical to each other in all of these organisms. These data suggest that observed differences between the regulatory tSH2 modules are the result of evolutionary selection and ancient specialization.

Using the MSA, the rate of sequence change at each position was calculated for both the Syk and ZAP-70 lineages. This rate is characterized by its Armon score; a lower score signifies a lower evolutionary rate, or a higher degree of conservation, at that position.^25^ In Figure 7, the Armon score is plotted for positions at which the human Syk and ZAP-70 residues have different properties (e.g. lysine and arginine are not considered to be different). Overall, the ZAP-70 tSH2 module is more conserved than that of Syk, with many residues in ZAP-70 having a low Armon score. However, there are positions at which the variance in ZAP-70 is high and the corresponding position in Syk is conserved, such as residues 246, 247, and 249. There are also many sequence differences that are conserved in both lineages. Notably many of these are located in the C-SH2 domain, which we have identified as a region where the two proteins show significant differences in dynamics.

**Figure 7.**
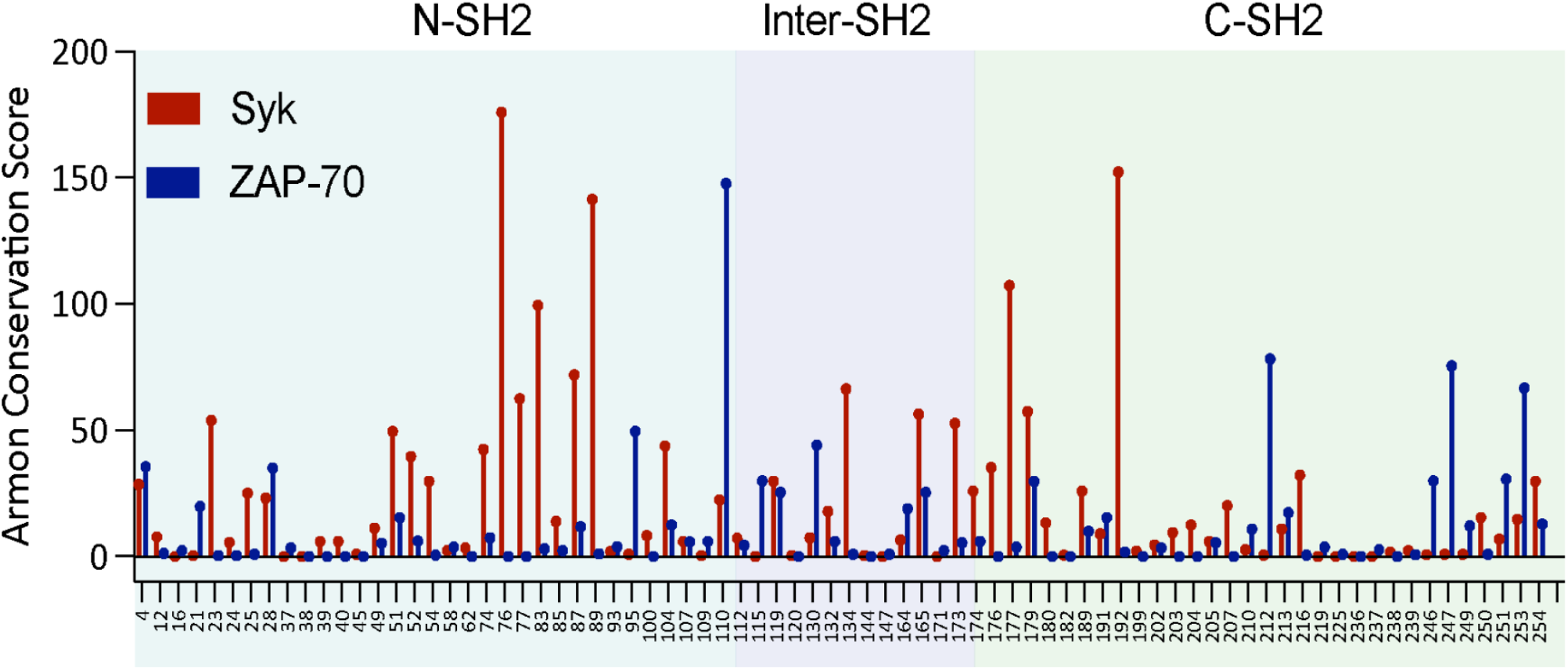
Sequence differences are conserved in both the Syk and ZAP-70 lineage. The Armon conservation score at each position (Syk numbering) where the human Syk (red) and ZAP-70 (blue) sequence differ significantly. Overall ZAP-70 is more conserved than Syk, but there are positions which are conserved in only the Syk lineage or equally conserved in both lineages.

A set of residues that are conserved in the Syk lineage but not in the ZAP-70 lineage was noted while considering the MD simulations. The root-mean-square-fluctuation (Å) (RMSF) of each Cα atom about its average position in the tSH2 modules is shown in Figure S1 (see supplementary information). The RMSF values calculated for residues ZAP-70 are higher than for Syk in several regions of the C-SH2. This region corresponds to the ends of three β strands as well as the loops connecting those strands (Figure 8A). Throughout the simulations two of these β strands (depicted in the lower portion of Figure 8B) frequently separate, and this separation is more sustained in the ZAP-70 simulations (Figure 8B and Figure S2). In the corresponding region of Syk we noted the presence of several charged amino acids, that are not present in ZAP-70 (Figure 8A). These charged amino acids may allow for the formation of cross β strand salt bridges, which stabilize a more closed conformation of these strands and contribute to the attenuated dynamics of the C-SH2 domain of Syk compared to those of the same domain in ZAP-70. Notably, these charged amino acids are highly conserved in Syk, but not in in ZAP-70 (Figure 8C), hinting at the specialization of not only the more globally conserved ZAP-70 but also Syk.

**Figure 8.**
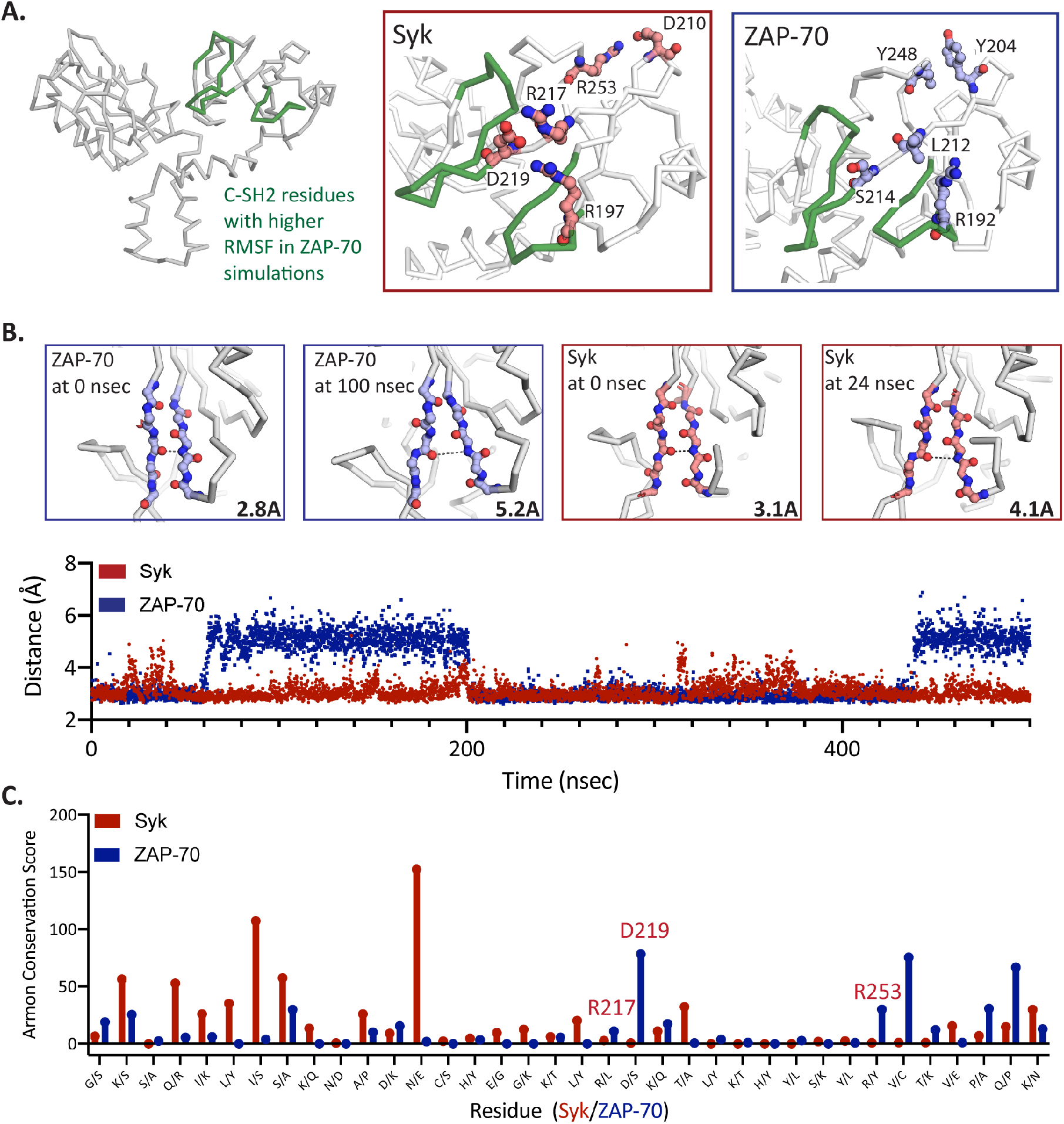
Charged residues conserved in Syk but not ZAP-70 may contribute to increased stability of the C-SH2. **A**. The residues in ZAP-70 with higher RMSF values than the corresponding regions of Syk are colored in green. Boxes highlight the C-SH2 domains of Syk (left) and ZAP-70 (right). In the Syk C-SH2 charged residues (red sticks) may be involved in stabilizing the fluctuations in this domain. D219 is able to interact with R217 and R197, forming cross B-strand interactions. These residues are not conserved in ZAP-70 (blue sticks). In Syk, R253 and D210 are positioned to interact. **B**. Snapshots of the B strands discussed in (A) at different points in the simulation. In both Syk and ZAP-70 these B-strands peel apart. The distance between two of the B-strand residues is noted in the corner of each box and is depicted by the dashed line. The lower panel shows this distance throughout one of the simulations (plots of other simulations can be found in Supplemental Figure 2). The B-strands in ZAP-70 move apart to a greater extent and this separation is sustained for longer than in Syk. **C**. All positions in the C-SH2 domain in which human ZAP-70 and human Syk differ are noted along the x-axis. Residues discussed in (A and B) are conserved in the Syk lineage but are not completely conserved in the ZAP-70 lineage, pointing to a potential constraint on this region during the evolution of Syk.

## DISCUSSION

We have determined the structure of the ITAM-free tSH2 module of Syk and found that its conformation differs from that of the ITAM-free tSH2 module of ZAP-70. The isolated ITAM-free Syk tSH2 module adopts a conformation similar to the ITAM-bound form, while the ZAP-70 tSH2 module adopts one similar to that seen in the structure of the full-length, autoinhibited kinase. Previously reported NMR residual dipolar coupling data for the ITAM-free Syk tSH2 module have shown that the SH2 domain-domain orientation observed in solution is similar to the orientation observed in the ITAM-bound state.^26^ This observation, as well as our HDX-MS data, indicate that our crystal structure reflects the solution structure of the Syk tSH2 module.

Klammt et al. previously used HDX-MS to probe full-length ZAP-70 and the isolated tSH2 module in various states representing steps of the activation pathway.^20^ Our HDX-MS data for the tSH2 module corresponds well to theirs; in particular, our conclusion that ZAP-70 is tuned towards an OFF state is consistent with earlier work. Recently, Gangopadhyay et al. proposed a model in which the C-SH2 domain of ZAP-70 binds the phosphorylated ITAM first with a low affinity, thus allowing the binding pocket of the N-SH2 to be fully formed.^19^ Binding of the second phosphorylated tyrosine in this pocket remodels the dynamics of the C-SH2 domain binding pocket, resulting in a higher affinity for the phosphorylated ITAM. We show that the C-SH2 of ZAP-70 is more flexible in the ITAM-free state and binding of a peptide rigidifies this domain. Significantly, we find that the same is not true for Syk, which is comparatively rigid, even in the ITAM-free state. We identified several, conserved polar residues in the C-SH2 of Syk which may contribute to stabilizing it in the absence of a bound phosphotyrosine, but additional studies are needed to elucidate how sequence differences contribute to the differences in conformation reported here.

Our evolutionary analysis of extant Syk and ZAP-70 homologs suggest that some sequence differences between the tSH2 modules are conserved in both the Syk and ZAP-70 lineages. We propose that the tSH2 module of Syk is less stable in its autoinhibited conformation, but the opposite is true for the tSH2 module of ZAP-70, which prefers the conformation observed in the auto-inhibited state. The additional conformational change required for activation in the tSH2 module of ZAP-70 likely provides another energetic barrier preventing spurious activation of ZAP-70 and thereby contributing to the tight control over the activation of T-cells. Other modes of regulation governing B-cell activation, such as the involvement of helper T-cells, may allow for Syk to be relatively less regulated, and could be important to its roles in B cells and other signal transduction pathways.

## MATERIALS AND METHODS

### Protein constructs and purifications

Both the Syk and ZAP-70 tSH2 modules were expressed and purified as described in Visperas et al.^27^

### Sequence Curation and Alignment

The ZAP-70 and Syk sequences were compiled through a series of BLAST searches, partially described in Shah et al.^28^ In total there are 184 sequences, 87 corresponding to ZAP-70, 89 to Syk, and 8 originating from hagfish and lamprey. Assignment as a Syk or ZAP-70 was determined based on highest homology to human Syk or ZAP-70. The sequences were aligned using the T-coffee sequence alignment software.^24^

### ITAM-free Syk crystallization, data collection, and structure determination

The Syk tSH2 was purified as described and then buffer exchanged via dialysis to a crystallography buffer containing 10mM HEPES pH 7.0, 50mM NaCl, 5% glycerol and 1mM TCEP. The dialyzed protein was immediately concentrated to 35mg/mL. 1ul of the concentrated protein was mixed with 1ul of well solution (0.2M sodium nitrate and 10% w/v PEG 3350) and equilibrated in a hanging drop set-up ∼20 °C. Crystals grew over the course of two weeks after which they were harvested. The crystals were soaked in a cryoprotectant containing 0.2M sodium nitrate, 20% PEG 3350, and 20% glycerol and flash cooled in liquid nitrogen. The data was collected at the Advanced Light Source (Lawrence Berkeley National Laboratory) on beamline ALS 8.2.2. Detailed data collection and structure determination procedures cane be found in the SI.

### HDX-MS

Deuterated buffer was prepared by three rounds of lyophilization of buffer (20 mM Tris pH 7.5, 150 mM NaCl, 0.5 mM TCEP, and 5% glycerol) followed by resuspension in D_2_O. ITAM-free experiments were carried out by diluting 30uM tSH2 module 10-fold into the deuterated buffer. At the following timepoints (15sec, 30sec, 1min, 5min, 15min, 30min, 1hour, and 4 hours) 80ul of the exchange protein was removed and quenched in a 5x quench buffer containing 4.8 M Gdn-Cl, 12% glycerol, and 2% formic acid. Quenched samples were immediately flash frozen in liquid nitrogen and stored at -80 °C. For the ITAM-bound experiments lyophilized peptide with the sequence corresponding to ITAM 1 of the T-cell antigen receptor ζ-chain (CGNQL(pY)NELNLGRREE(pY)DVLD; where pY is phosphotyrosine (prepared by David King, UC Berkeley/Howard Hughes Medical Institute) was resuspended in the same pH 7 buffer to a final concentration of 2.9mM. The concentrated peptide was mixed with 30uM protein to a final concentration of 260uM, well above the reported Kd of 80-100nM.^29^ This sample was allowed to equilibrate for four hours at room temperature. The equilibrated sample was then diluted 10x into deuterated buffer and timepoints collected and stored in the same way as the ITAM-free samples. Detailed descriptions of the LC-MS methods can be found in the SI.

### MD Simulations

For the MD simulations we began with the ITAM-bound tSH2 structures of Syk (PDB: 1A81 Chain A) and ZAP-70 (PDB: 2OQ1). To prepare these structures for simulation we edited the PDB files to remove the ITAMs as well as revert heavy atoms to their wildtype identity. We used the LEaP program in AMBER to solvate the proteins and add the counter ions (Na^+^ or Cl^-^) required to bring the net charge of the system to zero. For ZAP-70 the system size was 89,564 atoms and 82,514 atoms for Syk. For all simulations we used the AMBER ff14SB for protein atoms and the TIP3P for water and ion atoms.

All simulations were performed using the AMBER18 software and the Compute Unified Device Architecture (CUDA) version of Particle-Mesh Ewald Molecular Dynamics (PMEMD) on graphical processing units (GPU). The system energy was minimized in three rounds using the steepest-descent method and then heated to 300K. Heating was followed by 200psec of constant number/pressure/temperature equilibration. 500 or 1000nsec unconstrained simulations were started after energy minimization and equilibration, with coordinates saved every 2 ps. Replicate simulations were based on the same minimization model, but new equilibrations were performed in order to generate unique initial atom velocities. Additional methods in SI.

## Supporting information

Hobbs_et_al_SupplementaryMaterial

## ACKNOWLEDGEMENTS

We would like to thank the members of the Marqusee and Kuriyan labs for helpful discussions. We would also like to thank Emma Carroll and Naomi Latorraca for critically reading the manuscript. This work was supported by the US NIH R01GM050945 to SM, the US NIH PO1 5P01AI091580-10 to JK, and the Damon Runyon Cancer Research Foundation DRG-14 to NHS. SM is a Chan Zuckerberg Biohub Investigator. We also thank beamline staff from BL 8.2.2, part of the Berkeley Center for Structural Biology (BCSB) at the Advanced Light Source (ALS). The BCSB is supported in part by the Howard Hughes Medical Institute. The ALS is a Department of Energy Office of Science User Facility under Contract No. DE-AC02-05CH11231. The ALS-ENABLE beamlines are supported in part by the NIH, NIGMS, grant P30 GM124169.

## AUTHOR CONTRIBUTIONS

Conceptualization: HTH, JK, SM. Data curation: HTH and NHS. Investigation: HTH, JMB, and CLG. Visualization: HTH, NHS, JK, and SM. Writing: HTH, SM, and JK.

