## Supplementary material for "Differences in the dynamics of the tandem-SH2 modules of the Syk and ZAP-70 tyrosine kinases": Hobbs_et_al_SupplementaryMaterial

### SUPPLEMENTARY MATERIALS AND METHODS

*Structure determination of the isolated ITAM-free Syk tSH2 module*

Data was collected at the Advanced Light Source at the Lawrence Berkeley National Laboratory at $\lambda$= 0.999816 Å over 402°, with $\Delta\phi$=1° frames and an exposure time of 1 s per frame. The proper range for data collection was determined using iMOSFLM.[1] Data were processed using with XDS and then scaled and merged with Aimless in the CCP4 suite.[2–4] Molecular replacement was performed using Phenix Phaser using Chain F from the ITAM-bound structure as a search model (PDB: 1A18).[5,6] Refinement was performed using Phenix, manual refinement was done using Coot, and model geometry was assessed using Molprobity.[5–11] Coordinates and structure factors were deposited in the Protein Data Bank, with accession code .

*HDX-MS methods*

For both the apo and ITAM bound states, a tandem mass spec experiment was performed on undeuterated samples in order to sequence and identify peptides. Byonic (Protein Metrics) was used to generate peptide lists based on the tandem MS experiment. All samples were thawed immediately before injection into an LC system (Trajan and Thermo). The temperature of the LC columns was maintained at 4 °C in a cooled chamber in order to reduce back exchange. The quenched sample was subjected to an in-line digestion by two acid proteases, pepsin and fungal protease (Sigma). Following digestion, peptides were de-salted on a C4 trap column before analytical separation and elution with a 10-50% and then short 100% wash with 90% acetonitrile on a C8 analytical column. Peptides were eluted directly into a Q-Exactive Orbitrap mass spectrometer for analysis. Peptide deuteration states were determined by HD Examiner 3 (Sierra Analytics) by fits of the isotopic distribution to those predicted from the tandem mass spec experiment. Deuteration of peptides here is reported as #D incorporated, as calculated by HD Examiner 3.

*MD simulations-calculation of RMSF*

The AmberTools CPPPTRAJ package was used to analyze the simulations.[12] The distance between the SH2 domains was calculated as the distance between the center of mass of the N-terminal SH2 domain (residues 8-110) and the center of mass of the C-terminal SH2 domain (residues 161-252). We computed the root-mean-square fluctuation (RMSF) for each domain simulation by first determining its average structure across the simulation, after removing the first 30 nsec to ensure the simulation had sufficient time to equilibrate, and then calculating the RMSF from this average structure. The following residue numbers were used to align the Cα atoms of each domain in Syk: N-SH2 (9-110), C-SH2 (162-253), and inter-SH2 linker (111-161); and in ZAP-70: N-SH2 (9-110), C-SH2 (162-253), and inter-SH2 linker (111-161). The error bars depicted on the graphs of the RMSF refer to the standard deviation of RMSF values for each simulation, divided by the square root of the number of simulations per condition, or the standard error. Finally for the distance between the beta strands was calculated as the difference between the carbonyl oxygen of residue 212 and the backbone nitrogen of residue 197 for ZAP-70 and the carbonyl oxygen of residue 211 and the backbone nitrogen of residue 196 for Syk.

**Supplemental Table 1.** Data Collection and Refinement Statistics

| | **ITAM-free Syk tSH2 module** |
|---|---|
| **Wavelength** | |
| **Resolution range** | 45.78 - 3.2 (3.314 - 3.2) |
| **Space group** | C 1 2 1 |
| **Unit cell** | 143.882 153.881 85.472 90 91.049 90 |
| **Total reflections** | 61287 (6053) |
| **Unique reflections** | 30660 (3029) |
| **Multiplicity** | 2.0 (2.0) |
| **Completeness (%)** | 99.67 (99.11) |
| **Mean I/sigma(I)** | 7.02 (0.98) |
| **Wilson B-factor** | 73.77 |
| **R-merge** | 0.1197 (0.8567) |
| **R-meas** | 0.1693 (1.212) |
| **R-pim** | 0.1197 (0.8567) |
| **CC1/2** | 0.985 (0.367) |
| **CC*** | 0.996 (0.733) |
| **Reflections used in refinement** | 30597 (3018) |
| **Reflections used for R-free** | 1588 (167) |
| **R-work** | 0.2697 (0.3392) |
| **R-free** | 0.3039 (0.3440) |
| **CC(work)** | 0.916 (0.607) |
| **CC(free)** | 0.872 (0.682) |
| **Number of non-hydrogen atoms** | 11300 |
| **macromolecules** | 11296 |
| **solvent** | 4 |
| **Protein residues** | 1427 |
| **RMS(bonds)** | 0.002 |
| **RMS(angles)** | 0.67 |
| **Ramachandran favored (%)** | 95.66 |
| **Ramachandran allowed (%)** | 3.84 |
| **Ramachandran outliers (%)** | 0.50 |
| **Rotamer outliers (%)** | 9.04 |
| **Clashscore** | 9.07 |
| **Average B-factor** | 77.49 |
| **macromolecules** | 77.51 |
| **solvent** | 42.73 |

Statistics for the highest-resolution shell are shown in parentheses.

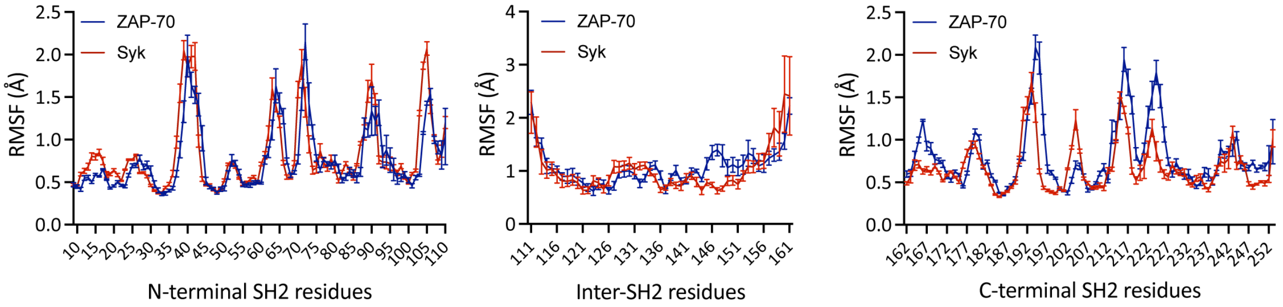


**Figure S1. The ITAM-free tSH2 module of ZAP-70 is more dynamic than that of Syk.** The C-SH2 domain of ZAP-70 and a section of the inter-SH2 linker are more dynamic than the corresponding regions in Syk as measured by the root-mean-square-fluctuation (Å) of each Cα atom in the individual domain about its average position in that domain across five independent simulations of each of the ITAM-free tSH2 modules. Error bars represent the SEM (n=5).

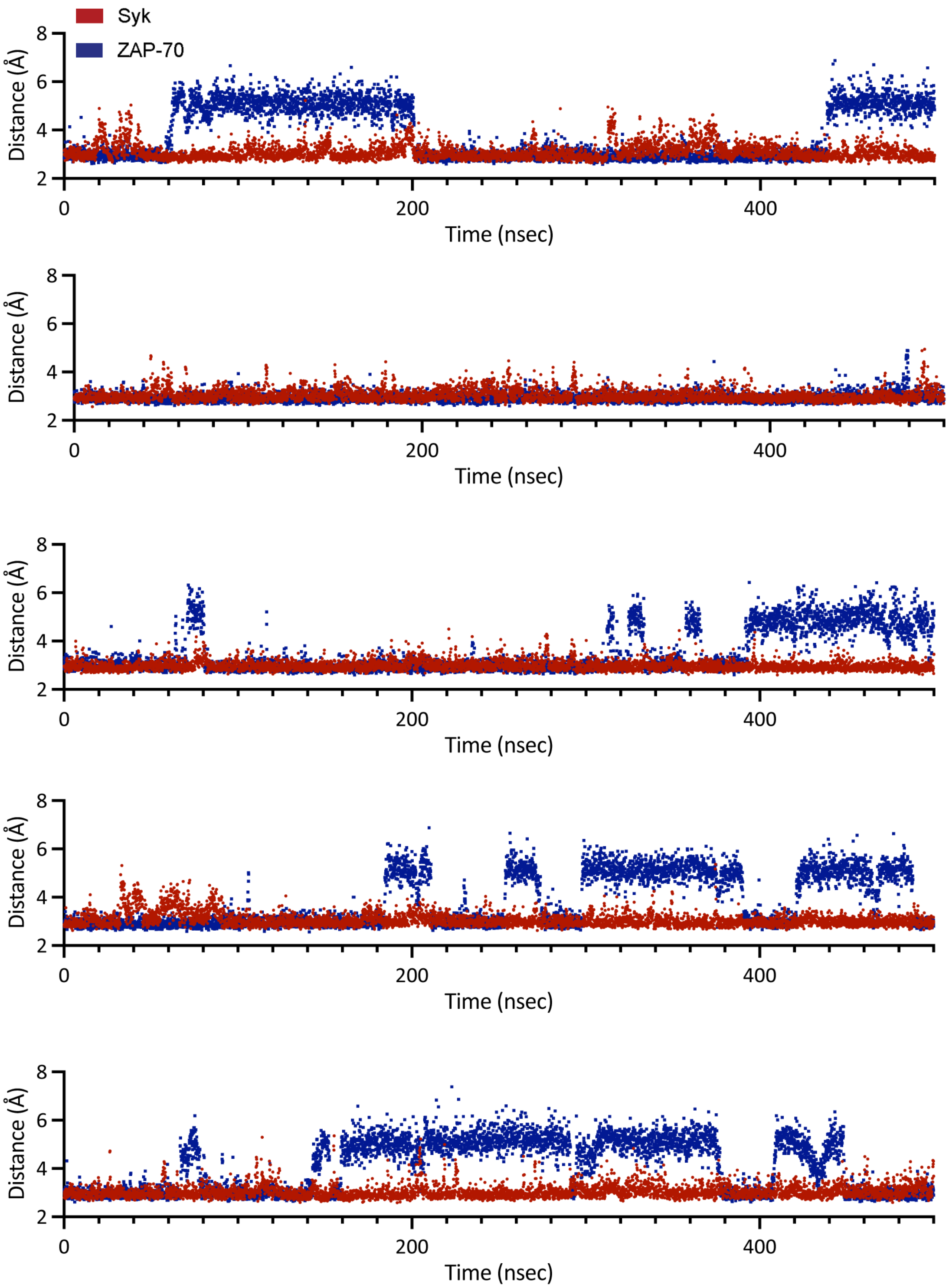


**Figure S2. Central β-strands in the C-SH2 of ZAP-70 fluctuate throughout the simulations.** The distance (Syk in red, ZAP-70 in blue) between two of the β-strand residues each pair of simulations. These β-strands in ZAP-70 move apart to a greater extent and this separation is sustained for longer than in Syk in most of the simulations.
